# Decreased integration of EEG source-space networks in disorders of consciousness

**DOI:** 10.1101/493395

**Authors:** Jennifer Rizkallah, Jitka Annen, Julien Modolo, Olivia Gosseries, Pascal Benquet, Sepehr Mortaheb, Hassan Amoud, Helena Cassol, Ahmad Mheich, Aurore Thibaut, Camille Chatelle, Mahmoud Hassan, Rajanikant Panda, Fabrice Wendling, Steven Laureys

**Affiliations:** Univ Rennes, LTSI, F-35000 Rennes, France; Azm Center for Research in Biotechnology and its Applications, EDST, Lebanese University, Lebanon; GIGA Consciousness, University of Liège, Liège, Belgium; Coma Science Group, University Hospital of Liège, Liège, Belgium

**Keywords:** Disorders of consciousness, high-density electroencephalography, functional brain networks, unresponsive wakefulness syndrome, minimally conscious state

## Abstract

Increasing evidence links disorders of consciousness (DOC) with disruptions in functional connectivity between distant brain areas. However, to which extent the balance of brain network segregation and integration is modified in DOC patients remains unclear. Using high-density electroencephalography (EEG), the objective of our study was to characterize the local and global topological changes of DOC patients’ functional brain networks.

Resting state high-density-EEG data were collected and analyzed from 82 participants: 61 DOC patients recovering from coma with various levels of consciousness (EMCS (n=6), MCS+ (n=29), MCS- (n=17) and UWS (n=9)), and 21 healthy subjects (i.e., controls). Functional brain networks in five different EEG frequency bands and the broadband signal were estimated using an EEG connectivity approach at the source level. Graph theory-based analyses were used to evaluate group differences between healthy volunteers and patient groups.

Results showed that networks in DOC patients are characterized by impaired global information processing (network integration) and increased local information processing (network segregation) as compared to controls. The large-scale functional brain networks had integration decreasing with lower level of consciousness.

## Introduction

Severe brain damages may lead to various disorders of consciousness (DOC; (Giacino *et al*., 2014)). Emerging evidence associates DOC with alterations in functional and/or structural brain networks, mainly those sustaining arousal and awareness (Owen *et al*., 2009; Boly *et al*., 2012; Fernández-Espejo *et al*., 2012; Annen *et al*., 2016; Amico *et al*., 2017; Bodien *et al*., 2017; Annen *et al*., 2018). Therefore, network-based EEG methods enabling the identification of these pathological alterations in brain networks are valuable. More specifically, new ‘neuromarkers’ able to identify network characteristics associated with DOC could improve diagnosis and optimize patient-specific clinical follow-up. This is important, since DOC encompass a spectrum from unresponsive wakefulness syndrome (UWS; wakefulness with only reflex movements) (Laureys *et al*., 2010; Monti *et al*., 2010), to minimally conscious state (MCS, reproducible and purposeful behavior; divided in MCS- and MCS+, the latter characterized by the presence of response to command, intentional communication and/or intelligible verbalization) (Giacino *et al*., 2002) and emergence from the minimally conscious state (EMCS, characterized by recovered functional communication and/or object use) (Giacino *et al*., 2002).

Electroencephalography (EEG) records cortical electrical activity from scalp electrodes, and has major assets due to its non-invasiveness, easiness-of-use and clinical accessibility. Previous EEG network-based studies in the context of DOC have been performed at the scalp level (Chennu *et al*., 2014; Chennu *et al*., 2017) with satisfactory accuracies in classifying UWS and MCS patients (Sitt *et al*., 2014; Chennu *et al*., 2017; Engemann *et al*., 2018). However, the biological interpretation of corresponding network alterations is not straightforward, since scalp EEG signals are corrupted by the volume conduction due to the head electrical conduction properties (Brunner *et al*., 2016; Van de Steen *et al*., 2016). Several studies have indeed reported the limitations of computing connectivity at the EEG scalp level (see for review (Schoffelen and Gross, 2009; Hassan and Wendling, 2018) even if this can be compensated by methods removing zero-lag components, (Vinck *et al*., 2011; Chennu *et al*., 2017). More essentially, scalp analysis does not allow making inferences about interacting brain regions. A potential solution is an emerging technique called “MEG/EEG source connectivity” (De Pasquale *et al*., 2010; Hipp *et al*., 2012; Mehrkanoon *et al*., 2014; Hassan *et al*., 2015; Kabbara *et al*., 2017; Mheich *et al*., 2017; Kabbara *et al*., 2018; Rizkallah *et al*., 2018), which reduces the aforementioned volume conduction. It is also conceptually attractive since networks can be directly identified at the cortical level with a high time/space resolution (for more details, see (Hassan and Wendling, 2018). Since conscious processing involves synchronization of locally generated oscillations between remote groups of neurons (Melloni *et al*., 2007), high-density EEG functional connectivity at the source level is a promising approach to track such synchronizations. In contrast, the time resolution of most fMRI techniques does not enable the detection of fast neural oscillations (e.g., 30-80 Hz range), which are involved in conscious perception and information transfer between regions (Fries, 2015), limiting the possibilities to study synchronization-based communication with fMRI. Over the past decade, graph theory has become a well-established approach in the field of network neuroscience (Fornito *et al*., 2016). It provides complementary information to source connectivity methods by quantifying functional and/or statistical aspects of identified brain networks. Among the few studies using graph theory in DOC, a common finding is the identification of disturbances in overall network integration, usually computed by modularity-based approaches (Crone *et al*., 2014; Demertzi *et al*., 2015; Chennu *et al*., 2017). However, to what extent the balance between EEG frequency-dependent network segregation (local information processing) and integration (global information processing) is altered in DOC remains elusive, which is the main objective of this paper. More specifically, large-scale communication (integration) between brain regions appears to involve rather low-frequency oscillations such as the theta rhythm, while local information processing rather involves high-frequency oscillations such as the gamma rhythm (Lisman and Jensen, 2013) for a detailed review). Here, we tackle the issue of the integration/segregation balance and its relationship with low/high frequency neuronal oscillations in patients with DOC. In this study, we combined EEG source connectivity with graph theory, applied to resting-state high-density-EEG (256 channels) data recorded from patients with DOC whose diagnosis has been established based on the Coma Recovery Scale-Revised (CRS-R; (Giacino *et al*., 2004)). Our specific objectives were to i) track alterations in functional connectivity of cortical brain networks according to consciousness levels (ranging from patients diagnosed as unresponsive, through those who have emerged from minimally conscious) and ii) identify the brain regions that were differentially involved between groups.

## Materials and Methods

### Participants

Sixty-one patients (24 females, mean age 40 ± 14.5) and twenty-one healthy subjects (i.e. controls; 8 females, mean age 41 years ± 15.4) were included in this study. Patients were diagnosed as EMCS (n=6), MCS+ (n=29), MCS- (n=17) and UWS (n=9). Etiology was traumatic in 28 patients and non-traumatic in 33 patients. Time since injury was on average three years and ranged from nine days to 19 years. The Ethics Committee of the University Hospital of Liège approved this study. All healthy subjects and patients’ legal surrogates gave informed written consent for participation to the study.

Patients’ level of consciousness was assessed using the CRS-R (Giacino *et al*., 2004) repeated at least 5 times to minimize clinical misdiagnosis (Wannez *et al*., 2017). Patient’s diagnosis was based on the best behaviors/highest item obtained over the repeated CRS-R assessments during the week of hospitalization. The following demographic information (listed in Supplementary Table T1) was also collected for each patient: age, gender, traumatic or non-traumatic etiologies and best clinical diagnosis based on the CRS-R assessments.

### Data acquisition and preprocessing

The full pipeline of the analysis is described in Figure 1. A high-density EEG system (EGI, Electrical Geodesic Inc., 256 electrodes applied with a saline solution) was used to record resting state brain activity with a sampling rate of either 250 Hz or 500 Hz (which were down-sampled to 250 Hz for consistency).

**Figure 1.**
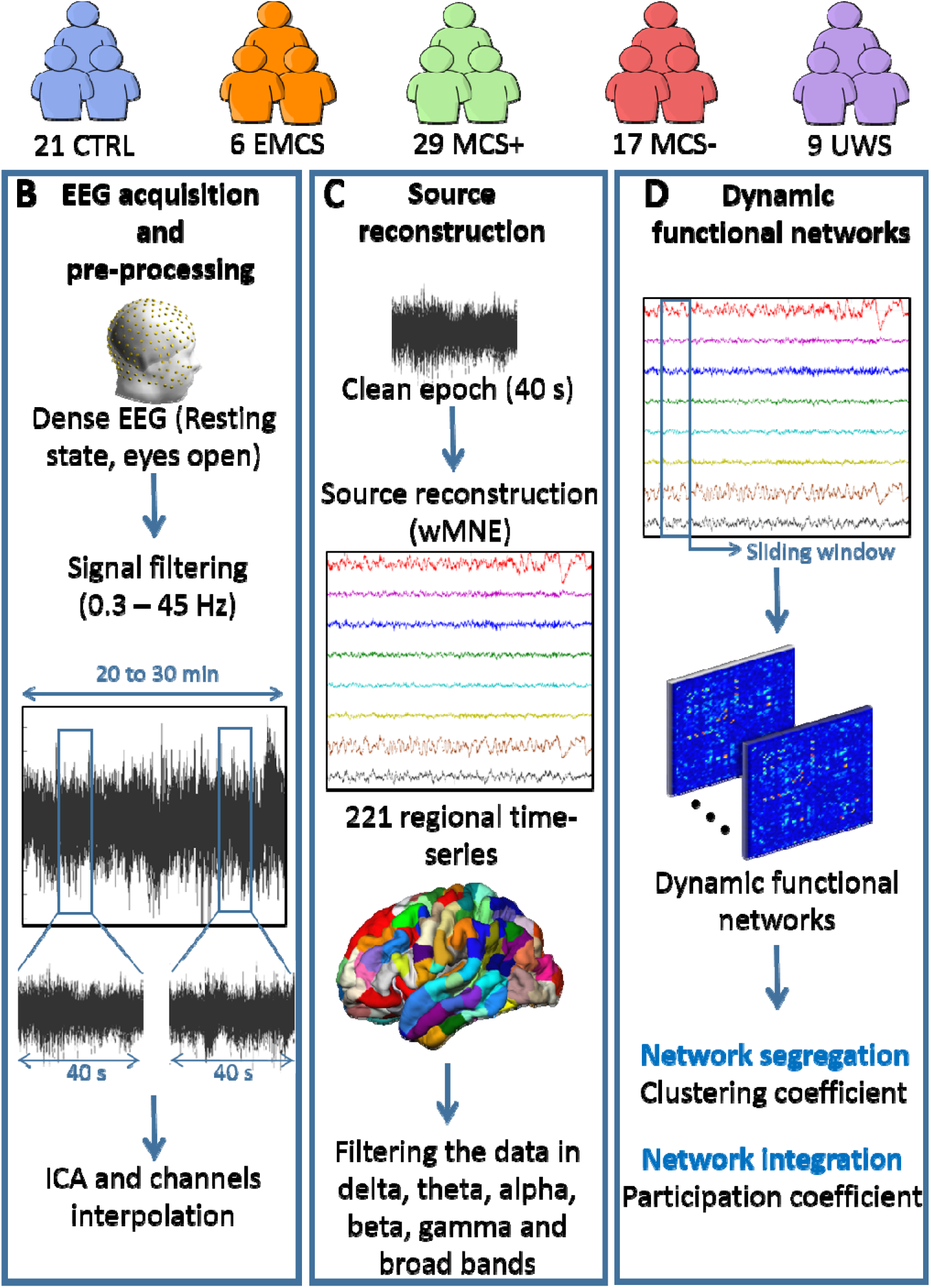
Data processing pipeline. (A) Database: Patients were diagnosed according to repeated assessments with the CRS-R into EMCS, MCS+, MCS- and UWS. The demographic details are listed in Supplementary Table T1. (B) EEG acquisition and preprocessing: High-density-EEGs were recorded using 256 electrodes during resting-state (eyes open, in the dark) for 20 to 30 min. Signals were then filtered between 0.3 and 45 Hz and segmented into 40 s epochs. Independent Component Analysis (ICA) was applied and bad channels were interpolated. Finally, the first five clean epochs were kept for analysis. (C) Source reconstruction: EEG cortical sources were estimated using the weighted norm estimation method (wMNE). This step was followed by a projection of the source signals on an atlas based on Desikan-killiany and Hagmann atlases, using a template brain. Reconstructed regional time series were filtered in six different frequency bands: Delta (1-3 Hz), Theta (3-7 Hz), Alpha (7-13 Hz), Beta (14-25 Hz), Gamma (30-45 Hz) and Broadband (1-45 Hz). (D) Dynamic functional networks: Functional connectivity matrices were computed using the phase locking value (PLV) calculated using a sliding window technique. Networks were then characterized by their clustering coefficient (segregation) and participation coefficient (integration).

EEG data from 178 channels on the scalp were retained for analysis; neck, forehead and cheeks channels were discarded, since they are the most prone to muscular artifacts, as previously described (Hassan *et al*., 2016; Kabbara *et al*., 2017). EEG signals were filtered between 0.3 and 45 Hz. Overall, out of 115 patients’ recordings, we retained 61 datasets for further processing and analysis. The other recordings were excluded due to excessive contamination by artefacts (e.g., muscle artefacts).

All EEG epochs were visually inspected before Independent Components Analysis (ICA) was performed to remove eye blinking artifacts using EEGLAB (Delorme and Makeig, 2004). Electrodes with poor signal quality were interpolated in Brainstorm (Tadel *et al*., 2011) using signals recorded by surrounding electrodes (spherical spline interpolating method, with a maximal distance between neighbors of 5 cm). The MRI template “Colin27” (Holmes *et al*., 1998) and EEG signals were co-registered through identification of the same anatomical landmarks (left and right tragus and nasion) using Brainstorm (without digitalizing the electrodes). The lead field matrix was then computed for a cortical mesh of 15000 vertices using openMEEG (Gramfort *et al*., 2010). The noise covariance matrix was calculated using a long segment of noisy EEG data at rest, as recommended in (Tadel *et al*., 2011). An atlas-based segmentation approach was used to project EEGs onto an anatomical framework consisting of 221 cortical regions identified by means of re-segmenting the Desikan-Killiany (Desikan *et al*., 2006) atlas using Freesurfer (Fischl, 2012). Time series within one region of interest were averaged after flipping the sign of sources with negative potentials.

### Brain networks construction

Functional brain networks were constructed using the “high-density-EEG source connectivity” method (Hassan *et al*., 2014) which quantifies the functional connectivity between regional time series at the source level. Reconstructed regional time series were filtered in six different frequency bands: Delta (1-3 Hz), Theta (3-7 Hz), Alpha (7-13 Hz), Beta (14-25 Hz), Gamma (30-45 Hz) and broadband (1-45 Hz). Then, we computed the functional connectivity between the reconstructed regional time series in each frequency band, using the phase locking value (PLV) (Lachaux *et al*., 1999). PLV values range between 0 (no phase locking) and 1 (full synchrony). Detailed methodological description and technical details of the high-density EEG source connectivity method as computed in this paper can be found in (Hassan *et al*., 2015).

We used a sliding window technique for each epoch to compute the dynamic functional connectivity matrices. The smallest window length recommended by (Lachaux *et al*., 2000) 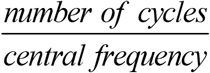 was used, equal to where the number of cycles at the given frequency band is equal to six. Finally, we adopted a 10% (of the highest PLV values) threshold to retain only the ‘true’ functional connections, and remaining PLV values were set to zero.

### Multi-slice networks modularity

Thresholded weighted connectivity matrices were split into time-varying modules using multi-slice networks modularity (Mucha *et al*., 2010; Bassett *et al*., 2013). This algorithm consists of linking nodes across network slices (time windows) *via* a coupling parameter before applying the modularity maximization method (Bassett *et al*., 2011; Bassett *et al*., 2015): each node is only connected to itself in the adjacent layers. This produces, for every brain region at every time window, a modular assignment reflecting the module allegiance.

Due to a degeneracy problem in the modularity algorithms (Good *et al*., 2010), i.e. running the same algorithm on the same connectivity matrix can result in slightly different outputs, the multilayer network modularity was computed 100 times and a 221 by 221 association matrix was generated (Lancichinetti and Fortunato, 2012; Bassett *et al*., 2013; Fornito *et al*., 2016). The association matrix elements indicate the number of times each node was assigned to the same module with the other nodes across these 100 partitions. The association matrix was then compared to a null-model generated from 100 random permutation from the originals partitions, and only significant values (*p* < 0.05) were kept (Bassett *et al*., 2011). Finally, the Louvain algorithm (Blondel *et al*., 2008) was applied on the association matrix to cluster the network, resulting in a partition that is the most representative of network modularity.

### Network measures

Our main intent was to explore two important properties related to information processing in the human brain network:

- *Network segregation*, which reflects local information processing. For this reason, the clustering coefficient ‘C’ was computed and considered as a direct measure of network segregation (Bullmore and Sporns, 2009). In brief, C represents how close a node’s neighbors tend to cluster together (Watts and Strogatz, 1998). This coefficient is the proportion of connections among a node’s neighbors, divided by the number of connections that could possibly exist between them, which is 0 if no connections exist and 1 if all neighbors are connected. The average clustering coefficient of a network was calculated for each epoch by averaging the clustering coefficient values over all the 221 regions.
- *Network integration*, which reflects global information processing. The participation coefficient was computed to measure the diversity of a node inter-modular connections (Guimera and Amaral, 2005). The participation coefficient of a node is close to 1 if its links are uniformly distributed among all the modules and 0 if all of its links are within its own module. Nodes with high participation coefficients interconnect multiple modules together, and hence can be seen connectivity hubs. The average participation coefficient of a network is calculated for each epoch by averaging the participation coefficient values over all 221 regions.

#### Statistical analysis

To statistically assess the difference in brain integration and segregation between healthy volunteers and all patient groups on the one hand, and between patient groups on the other hand, we used the Mann-Whitney U Test. In order to address the family-wise error rate, statistical tests were corrected for multiple comparisons using the Bonferroni method. In the global-wise analysis, *p*_bonf_ =0.05/N_g_, where N_g_ = 5 denotes the number of groups. In the region-wise analysis, *p*_bonf_ =0.05/N_r_, where N_r_=221 denotes the number of regions of interest. The Jonckheere-Terpstra (JT) test (Terpstra, 1952; Jonckheere, 1954), a non-parametric and rank-based trend test, was used to test the trends of network metrics as a function of the level of consciousness. Regional-level differences were analyzed between the groups, with the exception of the EMCS group due to its very small sample size (N=6).

## Data availability

The data used in the present study is available upon reasonable request.

## Results

There were no differences between patients and controls in terms of gender (*p*=0.6) or age (*p*=0.2) and between patient groups for time since injury (*p*=0.12).

### Network segregation/integration in DOC

The results in Figure 2A illustrate increased clustering coefficient values with decreased consciousness level in the delta (JT trend statistic=4.2, *p*<0.0001), theta (JT trend statistic=3.67, *p*=0.0001), beta (JT trend statistic=2.24, *p*=0.01) and gamma (JT trend statistic=1.66, *p*=0.04) bands. In the delta band, controls had lower clustering coefficients than patients in MCS+ (*p*=*0.0007, U=133, r=0.47, corrected*), MCS- (*p*<*0.0001, U=43, r=0.62, corrected*) and UWS (*p*=*0.01, U=39, r=0.485 corrected*). In the theta band, the clustering coefficient was also lower in controls as compared to MCS+ (*p*=*0.001, U=146, r=0.43, corrected*), MCS- (*p*=*0.0001, U=47, r=0.62, corrected*) and UWS (*p*=*0.01, U=38, r=0.46, corrected*).

**Figure 2.**
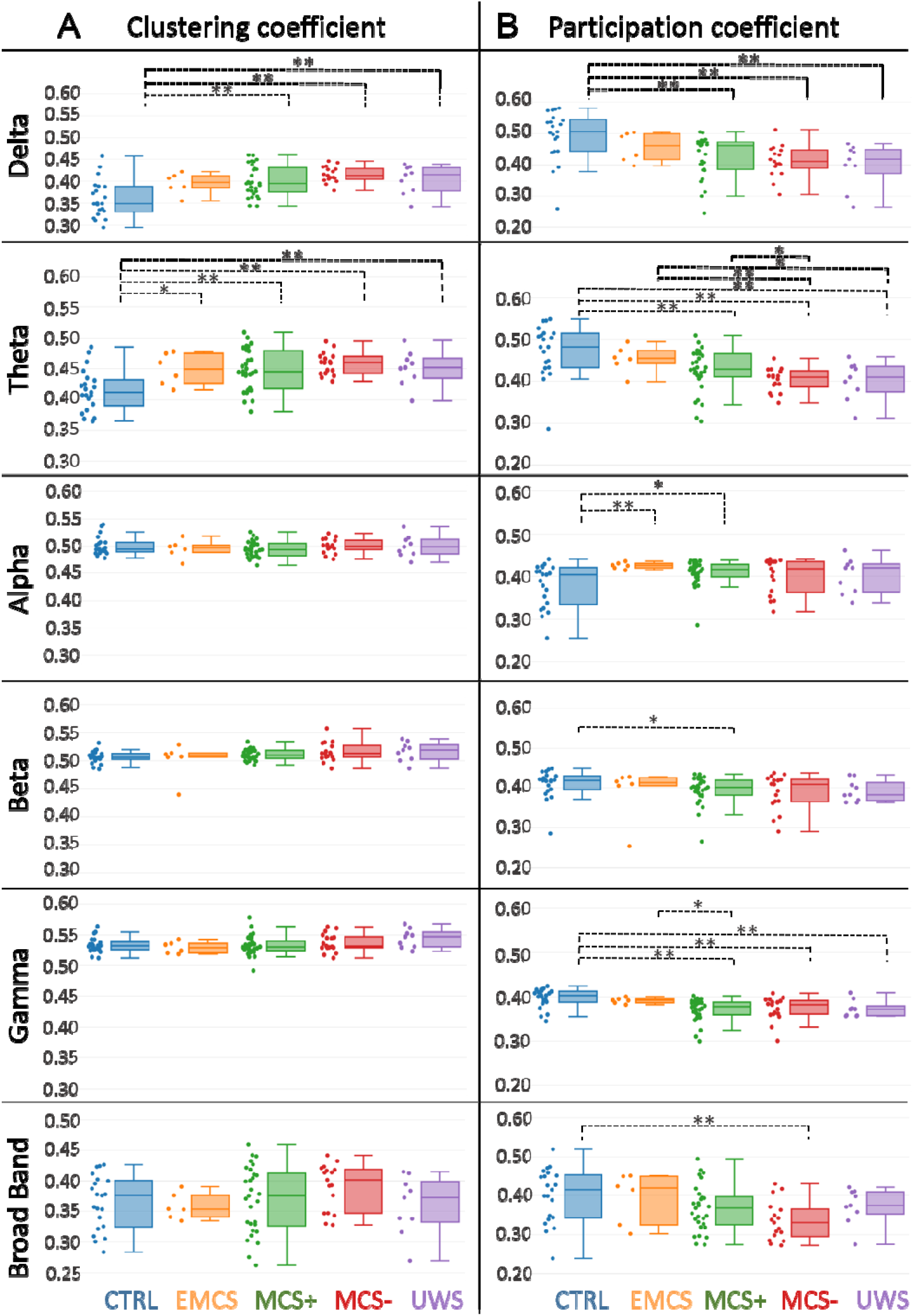
Brain segregation and integration in patients with disorders of consciousness. A. The clustering (segregation) and B. participation (integration) coefficients are presented for all groups in delta (1-3 Hz), theta (3-7 Hz), alpha (7-13 Hz), beta (14-25 Hz), gamma (30-45 Hz) and broad band (3-45 Hz). Values were averaged over all brain regions. Individual patient metrics are shown in the scatter plot next to the box plot. Increase of clustering coefficient values and decrease of participation coefficient values with decreased consciousness level was found within all frequency bands. A Wilcoxon test was applied between groups. * denotes *p*<*0.05* without correction and ** with correction.

The results in Figure 2B illustrate decreased participation coefficient values with decreased consciousness level in the delta (JT trend statistic=4.5, *p*<0.0001), theta (JT trend statistic=4.9, *p*<0.0001), beta (JT trend statistic=2.2, *p*=0.01), gamma (JT trend statistic=3.9, *p*<0.0001) and broad bands (JT trend statistic=2.7, *p*=0.0034). In the delta band, the participation coefficient was higher in controls as compared to MCS+ (*p*=*0.0005, U=480, r=0.48, corrected*), MCS- (*p*=*0.0007, U=294, r=0.54, corrected*) and UWS (*p*=*0.005, U=156, r=0.5 corrected*).

In the theta band, the participation coefficient was higher in controls as compared to MCS+ (*p*=*0.003, U=457, r=0.42, corrected*), MCS- (*p*<*0.0001, U=317, r=0.65, corrected*), and UWS (*p*=*0.003, U=159, r=0.52, corrected*). The participation coefficient in the theta band was also higher in EMCS as compared to MCS- (*p*=*0.007, U=90, r=0.56, corrected*).

In the alpha band, the participation coefficient showed lower values in controls than in EMCS (*p*=*0.009, U=18, r=0.49, corrected*). However, in the gamma band, the participation coefficient showed a decrease in MCS+ patients (*p*=*0.0002, U=492, r=0.52, corrected*), MCS-patients (*p*=*0.002, U=282, r=0.49, corrected*), and UWS patients (*p*=*0.01, U=150, r=0.45, corrected*) as compared to controls. Additionally, we performed the JT tests excluding the controls to confirm that our results were not solely driven by this group, and all trends remained significant in theta, delta and gamma bands.

### Regional-wise differences between groups

We present below results for the participation coefficient in the theta (Figure 3) and gamma (Figure 4) bands.

**Figure 3.**
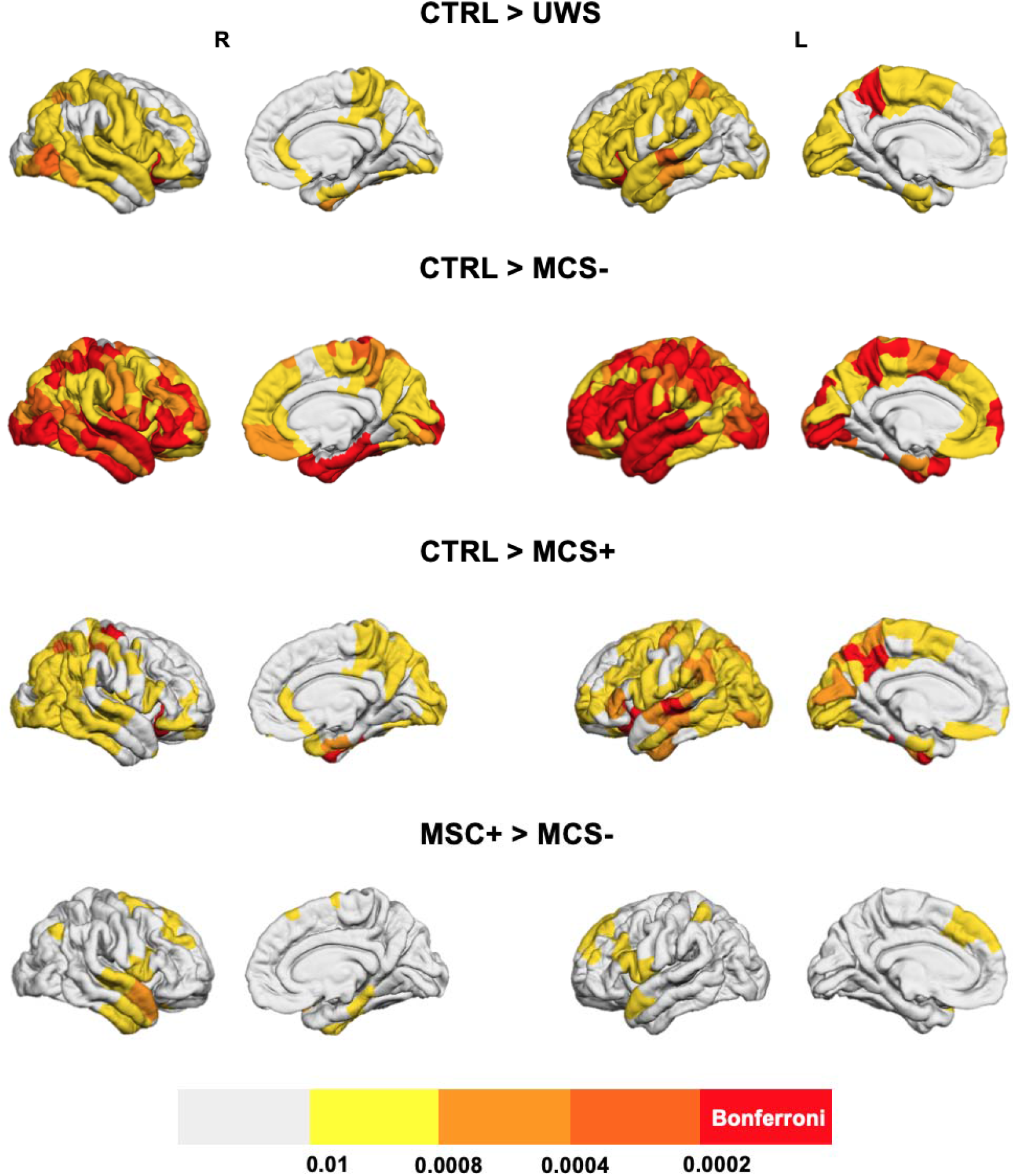
Between-group comparison of regional decreases in theta band integration. Brain regions that have significantly lower integration in UWS, MCS- and MCS+ as compared to the control group and in MCS-patients compared to MCS+ patients are presented. Brain regions having a p-value lower than 0.05/221=0.0002 (Bonferroni-corrected) are presented in the red color, regions with 0.0002<p<0.0004 are presented in dark orange, if 0.0004<p<0.0008 the light orange color was used, for 0.0008<p<0.01 the yellow color was used and if p>0.01 the regions are presented in white.

**Figure 4.**
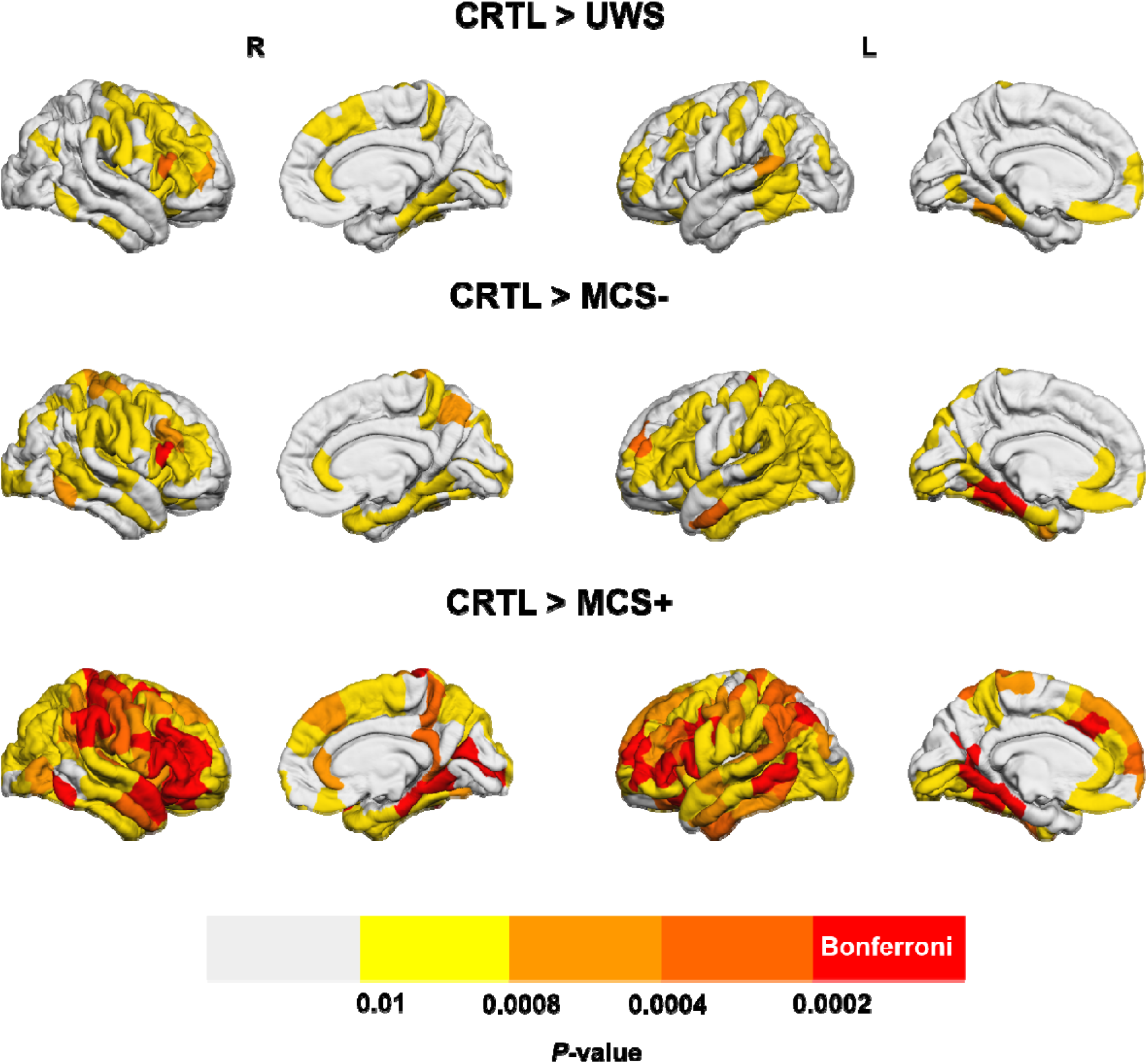
Between-group comparison of regional decreases in gamma band integration. Brain regions that have significantly lower integration in UWS, MCS- and MCS+ as compared to the control group are presented. Brain regions having a p-value lower than 0.05/221=0.0002 (Bonferroni-corrected) are presented in the red color, regions with 0.0002<p<0.0004 are presented in dark orange, if 0.0004<p<0.0008 the light orange color was used, for 0.0008<p<0.01 the yellow color was used and if p>0.01 the regions are presented in white.

Globally speaking, all patient groups had brain regions with significantly (Bonferroni-corrected) decreased integration as compared to the control group. As expected, a wider network was involved in the decreased integration in MCS-patients than in MCS+ patients, as compared to the controls. However, the differences between the control group and the UWS group were much less pronounced than between the control group and the MCS-group, which might originate from the small sample size of the UWS group (N=9). Regions that resisted the Bonferroni correction were mainly located in the left precuneus and left/right orbitofrontal area for the comparison of controls > UWS (*p*<0.0002). A large number of brain regions had decreased integration in the MCS-group as compared to the control group (exhaustive list in Supplementary Material), including the right orbitofrontal (*p*<0.0002,), left inferior temporal (*p*<0.0002) and left superior parietal (*p*<0.0002,). The left precuneus, left/right orbitofrontal, left/right fusiform, left superior temporal, right precentral (*p*<0.0002) showed a higher participation coefficient in the theta band for controls than MCS+ patients.

The results regarding the participation coefficient for the reconstructed functional networks in the gamma band are presented in Figure 4. No significant differences were observed between patient groups. The comparison between control and MCS-groups revealed a decrease in participation coefficient in MCS-patients mainly in the left fusiform, left postcentral and right dorso-lateral frontal cortex (*p*<0.0002). A much wider network of regions had a decreased participation coefficient between control and MCS+ groups, mainly located in the left/right lateral frontal cortex and right central cortex (*p*<0.0002). Again, as in the case of the theta band, the UWS group had a lower number of regions with decreased integration than MCS groups, as compared to the control group. The exact labels of the regions with a significant difference in the participation coefficient (in theta and gamma bands), along with Bonferroni corrections, are available in the Supplementary Materials Tables T2 and T3.

## Discussion

Emerging evidence supports that DOC are characterized by disruptions of brain networks that sustain arousal and awareness, as reviewed recently by (Bodien *et al*., 2017). Therefore, identifying alterations in whole-brain functional networks from non-invasive techniques, along with their relationships with varying consciousness levels, is a crucial and challenging issue. In this study, based on scalp high-density EEG recordings, we identified alterations in resting-state functional networks associated with DOC. Interestingly, a gradual reconfiguration of functional brain networks was observed in line with the consciousness level. Our findings also pointed at a decrease in brain network integration (communication between distant brain modules) and an increase in brain network segregation (communication within the same brain module) in DOC patients. A decrease in brain integration is especially relevant from a fundamental point of view. Indeed, one of the most prevalent theories, namely the Integrated Information Theory (IIT), describes the generation of conscious experiences as a result of a sufficiently complex integration of information between brain regions (Tononi, 2004; Tononi *et al*., 2016).

Although using EEG to identify markers in DOC is not novel in itself, the originality of the present work is that, as opposed to most previous studies, functional brain networks were estimated at the cortical level using high-density EEG data, which enabled making inferences about interacting regions. For example, we explored alterations of functional brain networks in two key aspects of human brain information processing: segregation and integration. These findings indicate that functional connectivity between distant areas (network integration) decreases with decreased level of consciousness.

### Low integration in DOC networks

The main finding of the present study is a decreasing trend in the integration of resting-state functional brain networks with the consciousness level. This finding is consistent with the conclusions from several studies (Crone *et al*., 2014; Chennu *et al*., 2017), including those investigating transcranial magnetic stimulation (TMS)-evoked EEG responses in patients with DOC (Casali *et al*., 2013; Casarotto *et al*., 2016). An important contribution of the present study is that, as opposed to complexity-based indexes computing using TMS-evoked responses, the computation of integration is based only on resting-state functional networks. It does not require any brain stimulation hardware, which could have a potential clinical and practical value, even if in this study we were not able to identify a difference between MCS and UWS groups, which remains a key clinical question.

More specifically, there were two common anatomical regions that were identified with decreased integration when comparing the control group with any of the patient groups. The first is the left precuneus, a key hub structure from the default mode network (DMN). The precuneus is engaged in self-related processing (Zhang and Chiang-shan, 2012), episodic memory (Ren *et al*., 2018), awareness and conscious information processing (Kjaer *et al*., 2001; Vogt and Laureys, 2005; Long *et al*., 2016). The second is a portion of the left orbitofrontal cortex, which is thought to encode predicted values of potential rewards (Gottfried *et al*., 2003). It is also though to play a major role in the evaluation of specific behavioral outcomes to influence action selection, depending on emotional and sensory contexts (Rudebeck and Murray, 2014). As mentioned previously, the network of regions with decreased integration in the theta band is wider in MCS-patients than in MCS+ patients, which is consistent with their respective clinical CRS-R diagnosis. It remains unclear however why the UWS group had fewer involved brain regions with decreased integration when compared to controls than the MCS+ and MCS-groups, which might be due to the small sample size of the UWS group (N=9).

Regarding the increased network segregation, results were less pronounced between groups as compared to network integration. This could explain discrepancies between previous studies, some referring to an increase in network segregation (using the clustering coefficient for instance) (Chennu *et al*., 2014), while others reported the opposite (Chennu *et al*., 2017). It is worth mentioning that, in our study and the one conducted by (Chennu *et al*., 2017), segregation results have opposite trends. A possible explanation is that the study by (Chennu *et al*., 2017) performed functional connectivity in the electrode space, while the present study functional connectivity was computed in the source space.

The relationship between scalp *versus* source space functional connectivity using EEG is indeed still an open question in the EEG community. A recent study that compared scalp- and source-reconstructed networks concluded that, not only the magnitude of network measures may change from scalp to source (EEG) analysis, but even the direction of the effect may be the opposite (still depending on the functional connectivity metric) between both methods (Lai *et al*., 2018). Therefore, even if further efforts need to be made in the comparison between source- and scalp-EEG-based networks, such discrepancies can be explained by the method of network reconstruction.

### Methodological considerations and limitations

In this study, a proportional threshold of 10% was used to eliminate spurious connections from connectivity matrices. We chose using a proportional threshold instead of an absolute threshold to warrant equal density between groups, as recommended by (van den Heuvel *et al*., 2017). Moreover, Garisson et al. (Garrison *et al*., 2015) reported that network measures are stable across proportional thresholds, as opposed to absolute thresholds. A variety of thresholding methods are available, but no method is free of bias, it is therefore recommended to perform studies across different values of thresholds to ensure that findings are robust against this methodological factor. Therefore, we tested several threshold values, which did not have any impact on the main conclusions of the study, as illustrated in Supplementary Figures S1 and S2.

A persistent problem in the field of MEG/EEG source functional connectivity is the volume conduction effect (Brookes *et al*., 2012). Source level connectivity analysis has been shown to diminish the volume conduction problem, since connectivity metrics are estimated between ‘local’ regional time-series. However, these ‘mixing effects’ can also arise in the cortical source space, and ghost couplings can be produced by some connectivity methods when applied to mixed signals. To tackle this issue, a number of methods were developed mainly centered on the zero-lag correlation rejection. Un-mixing methods, called ‘leakage correction’, have been reported to force the reconstructed signals to have zero cross-correlation at lag zero (Colclough *et al*., 2015). Although handling this problem -theoretically-improves interpretation, a recent study showed that current correction methods also produce erroneous human connectomes under very broad conditions (Pascual-Marqui *et al*., 2017).

One limitation of the present study is that a template source space was used, instead of a subject-specific one. This might be problematic in severely brain-injured patients, since different brain regions are injured between patients. In the case of healthy subjects, (Douw *et al*., 2018) found that co-registration with a template brain yielded largely consistent connectivity and network estimates as compared to native MRI. However, in the case of severe brain damage, it remains unknown how a template instead of a native MRI co-registration affects results and their interpretability.

Although the significant trends in network integration decrease with the level of consciousness, the current approach failed to identify a significant difference, at the group level, between MCS and UWS groups. This absence of difference represents, at this stage, a limitation in terms of potential clinical translation. That being said, this represents a challenge and an opportunity to develop further EEG network-based markers of the consciousness level based solely on resting-state recordings, to improve the diagnosis of DOC patients using a limited and accessible hardware.

## Supporting information

## Acknowledgements

This study was supported by the Future Emerging Technologies (H2020-FETOPEN-2014-2015-RIA under agreement No. 686764) as part of the European Union’s Horizon 2020 research and training program 2014–2018; the University and University Hospital of Liège; the Belgian National Funds for Scientific Research (FRS-FNRS); the Human Brain Project (EU-H2020-fetflagship-hbp-sga1-ga720270); the French Speaking Community Concerted Research Action (ARC - 06/11 - 340); NSERC discovery grant, IAP research network P7/06 of the Belgian Government (Belgian Science Policy); the European Commission; the James McDonnell Foundation; Mind Science Foundation; the BIAL foundation; the European space agency (ESA); the Public Utility Foundation ‘Université Européenne du Travail’. JR is a PhD student with a fellowship from the AZM and SAADE Association, Tripoli, Lebanon. OG is postdoctoral researcher at FRS-FNRS. OG is post-doctoral fellow, and SL is research director at the F.R.S.-FNRS. CC is a post-doctoral Marie Skłodowska-Curie fellow (H2020-MSCA-IF-2016-ADOC-752686).

## Declaration of authorship

Conceptualization: PB, MH, FW, SL. Data curation: JA, OG, SM, HC, AT, CC, RP, SL. Formal analysis: JR, JM, PB, MH, FW. Funding acquisition: FW, SL. Investigation: JR, JA, JM, PB, MH. Methodology: JA, PB, MH, FW, SL. Project administration: FW, SL. Resources: FW, SL. Software: JR, MH. Supervision: FW, SL. Validation: FW, SL. Visualization: JR, JA, JM, MH. Writing (original draft): JR. Writing (review and editing): JR, JA, JM, OG, PB, SM, HA, HC, AM, AT, CC, MH, RP, FW, SL.

