## Supplementary material for "Decreased integration of EEG source-space networks in disorders of consciousness"

| **Name** | **Age** | **Gender** | **Days since injury** | **Etiology** | **Best diagnosis** |
| --- | --- | --- | --- | --- | --- |
| P1 | 27 | F | 1570 | NT | MCS+ |
| P2 | 27 | M | 1542 | T | MCS+ |
| P3 | 35 | M | 6950 | NT | UWS |
| P4 | 60 | M | 9 | NT | MCS- |
| P5 | 24 | M | 319 | T | MCS- |
| P6 | 30 | F | 2406 | NT | MCS- |
| P7 | 30 | F | 563 | T | MCS- |
| P8 | 30 | M | 583 | T | MCS+ |
| P9 | 50 | M | - | T | MCS+ |
| P10 | 30 | F | - | T | MCS+ |
| P11 | 46 | M | 528 | T | MCS+ |
| P12 | 48 | F | - | NT | MCS- |
| P13 | 37 | M | 1869 | NT | MCS- |
| P14 | 59 | F | - | NT | MCS- |
| P15 | 5 | F | - | T | MCS+ |
| P16 | 24 | M | 2681 | NT | MCS+ |
| P17 | 30 | M | 33 | NT | MCS+ |
| P18 | 43 | M | 3139 | T | MCS- |
| P19 | 45 | F | 491 | NT | UWS |
| P20 | 57 | M | 390 | NT | MCS+ |
| P21 | 25 | F | 308 | NT | MCS+ |
| P22 | 23 | M | 421 | T | MCS+ |
| P23 | 28 | M | 66 | NT | MCS- |
| P24 | 53 | M | 1235 | NT | MCS- |
| P25 | 24 | M | - | T | MCS+ |
| P26 | 36 | F | - | NT | UWS |
| P27 | 22 | M | 2972 | T | MCS- |
| P28 | 23 | M | 2035 | T | MCS+ |
| P29 | 73 | M | 28 | NT | MCS- |
| P30 | 30 | M | 3337 | T | MCS+ |
| P31 | 47 | F | - | NT | MCS- |
| P32 | 65 | M | 674 | T | MCS+ |
| P33 | 55 | M | - | NT | MCS+ |
| P34 | 19 | M | 426 | T | MCS+ |
| P35 | 39 | F | 1437 | T | MCS- |
| P36 | 34 | F | 375 | T | EMCS |
| P37 | 61 | F | 858 | NT | MCS+ |
| P38 | 14 | M | 185 | NT | EMCS |
| P39 | 26 | F | 112 | NT | UWS |
| P40 | 35 | M | 4154 | NT | MCS+ |
| P41 | 60 | M | 406 | NT | EMCS |
| P42 | 62 | M | 672 | NT | UWS |
| P43 | 67 | F | 1464 | NT | MCS+ |
| P44 | 23 | M | 456 | NT | UWS |
| P45 | 42 | F | 220 | NT | MCS- |
| P46 | 72 | M | 3062 | NT | MCS+ |
| P47 | 21 | M | 257 | T | UWS |
| P48 | 30 | M | 402 | T | MCS- |
| P49 | 28 | M | 2423 | T | EMCS |
| P50 | 59 | F | 709 | T | MCS+ |
| P51 | 51 | F | 347 | NT | UWS |
| P52 | 25 | M | 1283 | T | MCS+ |
| P53 | 42 | M | 1186 | T | EMCS |
| P54 | 24 | F | 333 | NT | MCS- |
| P55 | 43 | F | 40 | T | UWS |
| P56 | 55 | F | 669 | T | MCS+ |
| P57 | 54 | M | 387 | NT | MCS+ |
| P58 | 38 | M | 541 | T | MCS+ |
| P59 | 43 | F | 98 | NT | MCS+ |
| P60 | 22 | M | 423 | T | MCS+ |
| P61 | 33 | F | 308 | NT | EMCS |

**Table T1: Patients demographic information: age, gender, days since injury, traumatic (T) or non-traumatic (NT) etiology and clinical best diagnosis.**

| CTRL>UWS | CTRL>MCS- | CTRL>MCS+ | MCS+>MCS- |
| --- | --- | --- | --- |
| Lateralorbitofrontal R  Lateralorbitofrontal L  Precuneus L | **Lateralorbitofrontal R**  **Lateralorbitofrontal L**  **Postcentral R**  **Inferiortemporal R**  **Parahippocampal R**  **Fusiform R Superiortemporal R**  **Superiortemporal L**  **Precentral R**  **Lateraloccipital R**  **Lateraloccipital L**  **Parsobitalis R**  **Rostralmiddlefrontal R**  **Rostralmiddlefrontal L**  **Inferiorparietal R**  **Bankssts R**  **Superiorparietal R**  **Middletemporal R**  **Middletemporal L**  **Supramarginal R**  **Temporalpole R**  **Parsopercularis R**  **Parsopercularis L**  **Pericalcarine R**  **Entorhinal R**  **Postriangularis R**  **Caudalmiddlefrontal L**  **Transvertemporal L** | **Fusiform R**  **Fusiform L**  **Lateralorbitofrontal R**  **Lateralorbitofrontal L**  **Precentral R**  **Precuneus L**  **Insula L**  **Superiortemporal L** |  |

**Table T2: Brain regions that have significantly lower integration in theta band in UWS, MCS- and MCS+ as compared to the control group and in MCS- patients compared to MCS+ patients with p-value lower than 0.05/221=0.0002 (Bonferroni-corrected)**

| CTRL>UWS | CTRL>MCS- | CTRL>MCS+ |
| --- | --- | --- |
|  | **Parsopercularis R**  **Fusiform L**  **Parahippocampal L**  **Lingual L**  **Postcentral L** | **Parsopercularis R**  **Rostralmiddlefrontal R**  **Rostralmiddlefrontal L**  **Parstriangularis R**  **Parstriangularis L**  **Lateralorbitofrontal R**  **Lateralorbitofrontal L**  **Parsopercularis R**  **Postcentral R**  **Postcentral L**  **Parsobitalis R**  **Superiortemporal R**  **Lingual R**  **Lingual L**  **Inferiortemporal R**  **Precentral R**  **Precentral L**  **Pericalcarine**  **Superiorfrontal R**  **Superiorfrontal L**  **Supramarginal R**  **Parahippocampal R**  **Parahippocampal L**  **Fusiform L**  **Caudalmiddlefrontal L**  **Precuneus L**  **Middletemporal L**  **Parsopercularis L**  **Inferiorparietal L** |

**Table T3: Brain regions that have significantly lower integration in gamma band in UWS, MCS- and MCS+ as compared to the control group and in MCS- patients compared to MCS+ patients with p-value lower than 0.05/221=0.0002 (Bonferroni-corrected)**

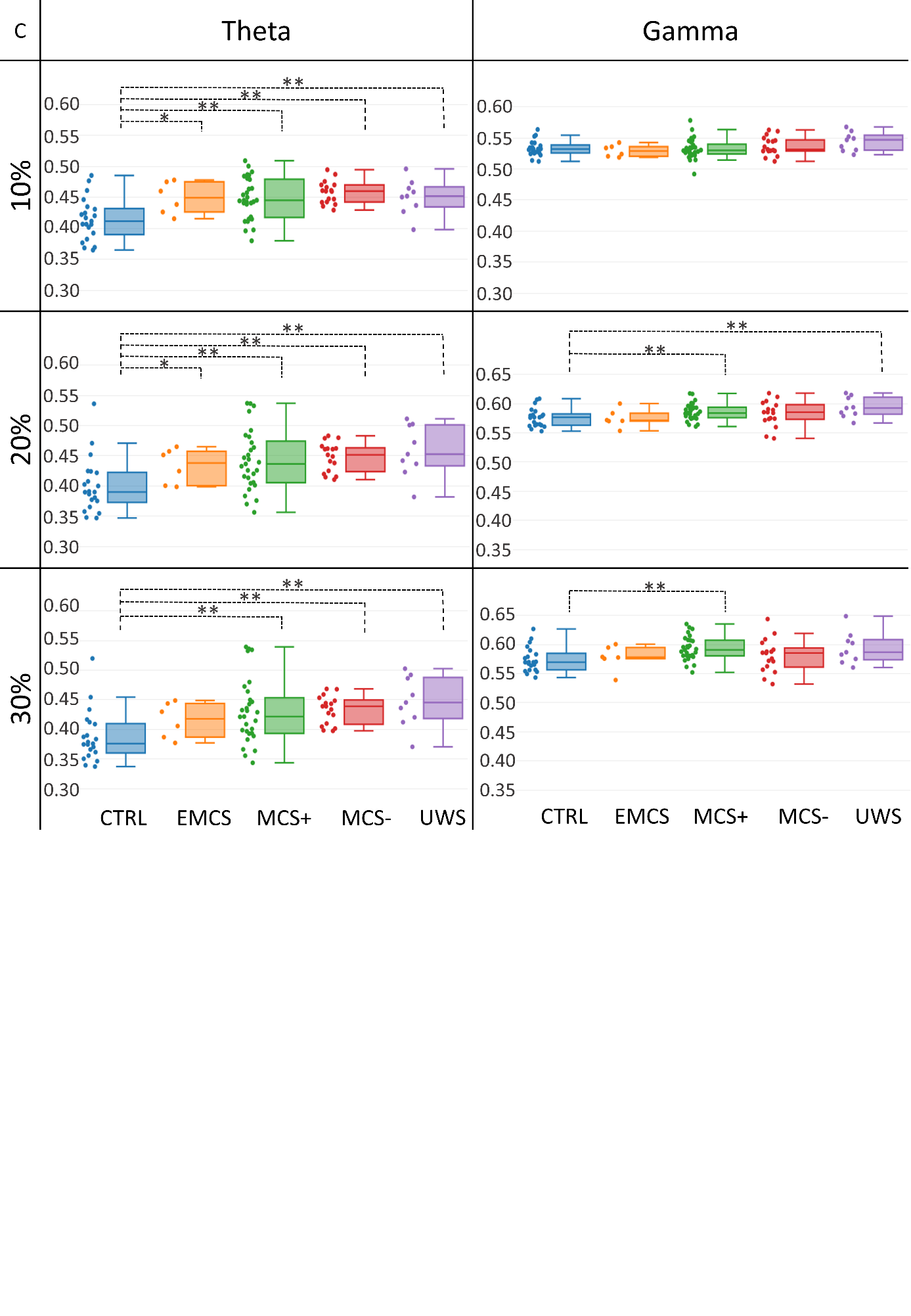

**Figure S1: Clustering coefficient values for different thresholds (10%, 20% and 30%) in the theta and gamma bands. Significant differences are presented by * if p<0.05 without correction and ** if corrected (Bonferroni correction p<0.05/5).**

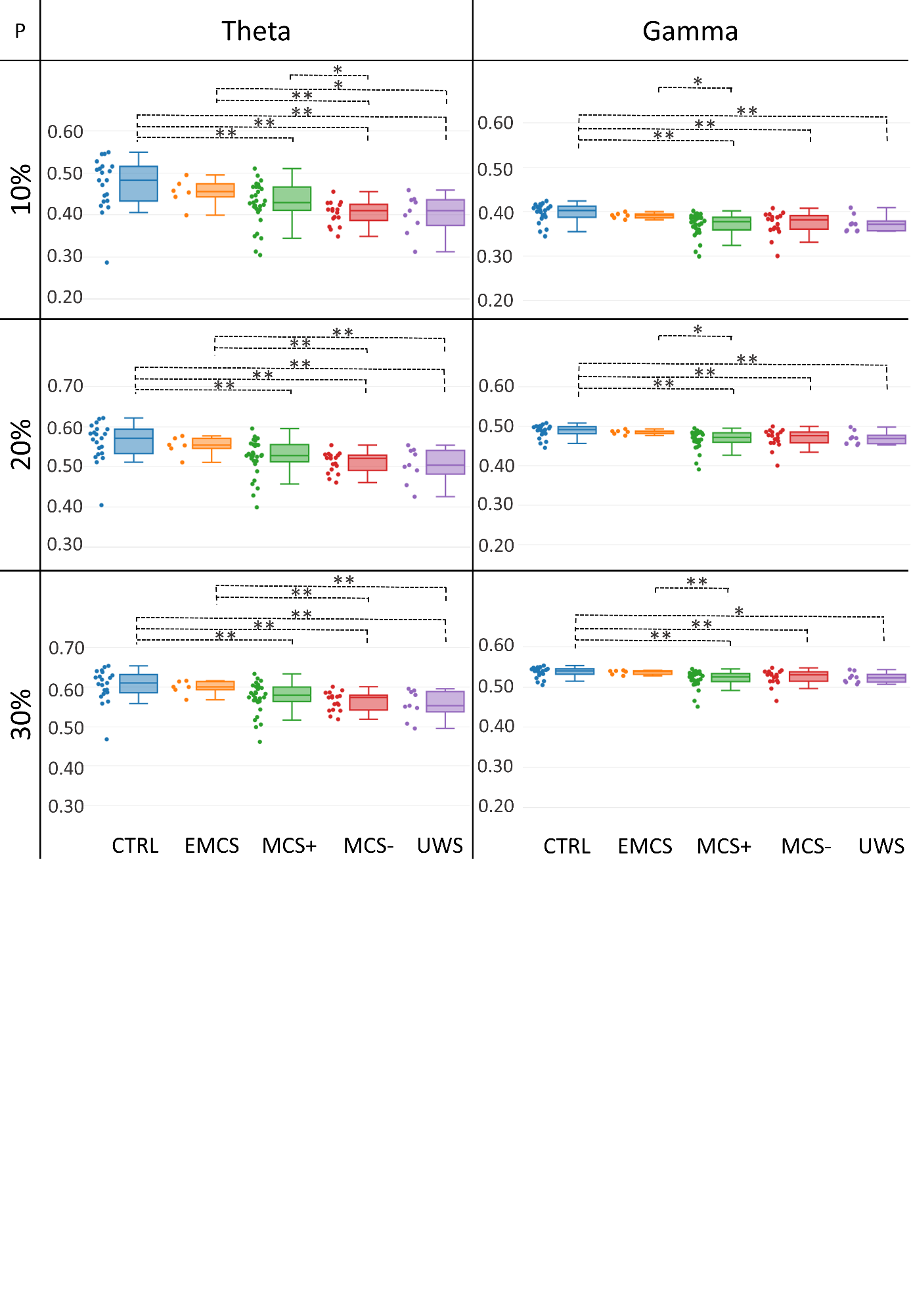

**Figure S2: Participation coefficient values for different thresholds (10%, 20% and 30%) in the theta and gamma bands. Significant differences are presented by * if p<0.05 without correction and ** if corrected (Bonferroni correction p<0.05/5)**
